# Optineurin is a gatekeeper of mitochondrial health and proteostasis in Alzheimer’s disease vulnerable neurons

**DOI:** 10.64898/2026.03.15.711617

**Authors:** Christina Tsagkogianni, Matteo Trivisonno, Johanna Sophie Willner, Carlos Garcia-Molinero, Yingyu Tang, Beatrice Mattina, Maria Latorre-Leal, Wei Wang, Jean-Pierre Roussarie, Patricia Rodriguez-Rodriguez

## Abstract

Alterations in autophagy-related pathways and in mitochondrial function have long been associated with the pathology of several neurodegenerative disorders, including Alzheimer’s disease (AD). However, the cascade of events that links these processes and how they contribute to the early degeneration of specific neuronal subpopulations remain to be understood. Here, we use a data-driven approach and identify *Optn* as a potential regulator of AD pathology that is highly enriched in vulnerable ECII neurons compared to neurons that degenerate later in the disease continuum. We show that Optineurin downregulation triggers early dysregulation of mitochondrial function, followed by alterations in AD-associated processes, including proteostasis, synaptic function, and neuroinflammation. This is accompanied by ECII neuron loss and astrocyte reactivity in EC neuron projecting areas in the hippocampus. Together our results suggest that Optineurin plays a central role in the maintenance of mitochondrial health and bioenergetics in AD vulnerable neurons and that pathological processes that impair this homeostasis may contribute to the early degeneration of vulnerable ECII neurons.

## Introduction

Neurodegenerative disorders are characterized by the accumulation of protein aggregates in specific brain regions and cell subpopulations within those regions, ultimately leading to neurodegeneration. In the case of Alzheimer’s disease (AD), those protein aggregates consist of extracellular amyloid beta peptides, known as amyloid plaques, and intraneuronal aggregates of hyperphosphorylated tau protein (neurofibrillary tangles, NFTs). Interestingly, and unlike other neurodegenerative disorders, these two pathological hallmarks do not follow the same deposition pattern within the brain: at prodromal disease stages amyloid plaques are found in the neocortex and hippocampus, while NFTs are limited to principal neurons located in layer II of the entorhinal cortex (ECII)^1-3^. Amyloid pathology is considered a major driver of the disease, a notion that is further supported by early-onset AD cases caused by mutations that result in enhanced pathological processing of amyloid precursor protein (APP). However, NFTs show a better correlation with clinical disease symptoms and with regional patterns of atrophy^4-6^. While ECII neurons have long been known to degenerate at very early disease stages^7, 8^, the reasons behind the selective vulnerability of ECII neurons to tau pathology and degeneration as well as the mechanisms that govern the interplay between amyloid and tau buildup in AD remain to be understood.

To shed light into the molecular mechanisms driving selective neuronal vulnerability in AD, we combined molecular signatures of ECII neurons with genomics data and built an in-silico model of gene networks within ECII neurons. Combining this network model with genome-wide association study data for NFT formation^9^ by using the NetWAS 2.0 (<u>Net</u>work-<u>W</u>ide <u>A</u>ssociation <u>S</u>tudy 2.0) machine-learning based algorithm, we re-prioritized genes based on their association with human tau pathology in vulnerable neurons. This approach identified four functional modules predicted to contribute to NFT formation in AD^10^. Genes within two of these modules were affected by amyloid pathology, what suggests that they may represent processes that interconnect these two pathological hallmarks.

In the present work we focus on the top predicted regulatory genes for each of these modules and combine this information with translatome profiling data from ECII neurons and neurons that are more resistant to AD. We identify *Optn* as a gene that is both highly enriched in ECII neurons compared to other neurons that are more resistant to AD pathology and predicted to have an association with ECII neuron vulnerability by NetWAS 2.0. By modulating *Optn* expression *in vivo* and *in vitro*, we identify Optineurin as a central regulator of mitochondrial health in AD vulnerable neurons. Optineurin downregulation triggers alterations in mitochondrial function that progressively lead to impaired synaptic function, proteostasis-related pathways, ECII neuron death and neuroinflammation characterized by astrocyte reactivity in EC neuron projecting regions in the hippocampus. These results suggest that mitochondrial homeostasis is especially relevant for AD vulnerable neurons, and highlight Optineurin as a potential therapeutic target in AD.

## Results

### *OPTN* is an ECII neuron-enriched predicted driver of tau pathology

As mentioned before, in previous work we developed a data-driven method to predict drivers of tau pathology in AD vulnerable neurons^10^. This approach identified four modules associated with tau pathology in AD vulnerable neurons. In the present work we aimed to identify genes that are enriched in ECII neurons and that are also top regulators of those four tau-pathology associated modules. We selected the five genes with highest connectivity scores to each of the modules, this is, genes that show a strong functional association as module regulators (**Figure S1A**). To identify differences in the biology of neuron subpopulations that may drive their resistance or vulnerability to AD neuropathology, we had previously generated profiles of actively translated mRNA in neurons in hippocampal CA1, CA2, CA3, dentate gyrus, somatosensory cortex and visual cortex or to ECII neurons, respectively^10^ using bacTRAP (<u>b</u>acterial <u>a</u>rtificial <u>c</u>hromosome – <u>T</u>ranslating <u>R</u>ibosome <u>A</u>ffinity <u>P</u>urification). We explored this dataset and found that, within the top module-connected genes, both *Optn* (connected to module 0) and *Piezo1* (connected to module 1) are highly enriched in ECII neurons compared to neurons that are more resilient to the disease (FDR = 6.68E-25 and 1.46E-8, respectively; **Figure S1A**). Considering that *Piezo1* showed very low expression levels in ECII neurons and is less consistently enriched in ECII neurons compared to all other cell types (**Figure S1B**), we decided to focus on *OPTN*.

*OPTN* codes for Optineurin, an autophagy adaptor that recognizes ubiquitinated cargo and has been shown to participate in the degradation or different cargos, including dysfunctional mitochondria^11, 12^ and pathological tau^13, 14^. Interestingly, several polymorphisms in *OPTN* have been associated with amyotrophic lateral sclerosis (ALS) and normal tension glaucoma, both characterized by axon degeneration^15, 16^.

### *Optn* downregulation in ECII neurons *in vivo* triggers DNA damage and neuroinflammation

To characterize optineurin function in ECII neurons *in vivo*, we overexpressed or silenced *Optn* by adeno-associated virus (AAVs) transduction in ECII-neuron bacTRAP mice (**Figure 1A)**. This model allows the analysis of gene expression changes triggered by *Optn* modulation specifically in ECII-neurons. We found no gene with significant differential expression in ECII neurons 3 weeks after *Optn* overexpression (log2FC = 1.12, FDR = 1.06E-5), suggesting that the levels of *Optn* at baseline might not be limiting for the proper function of ECII neurons, and that increasing its expression has no deleterious effects. On the other hand, *Optn* silencing (log2FC = -1.03, FDR = 0.014) led to widespread gene expression changes in these neurons 3 weeks after transduction (**Figure 1A-B, Supplementary Table 1**). Pathway analysis (Ingenuity Pathway Analysis, IPA) showed that some of the most significantly altered pathways in *Optn*-silenced neurons included several pathways involved in neuroinflammation (P = 1.07E-03) and DNA damage response, such as p53 (P = 2.95E-04) and ATM signaling (P= 4.17E-04). Axonal guidance signaling was among the most significant altered pathways and showed the highest number of associated genes (P = 3.02E-03) (**Figure 1C**). To further gain insight into alterations driven by Optineurin deficiency and in a relevant biological context, we used the functional module detection tool from the HumanBase resource^17^. HumanBase uses a given gene list (in this case DEGs upon *OPTN* silencing in ECII-neurons *in vivo*) and groups them into functional modules based on specific cell-types or tissues, in our case neurons. This method identified 3 different functional modules altered in *Optn*-silenced neurons (**Figure 1D**). In agreement with IPA analysis, one of these functional modules was enriched in DNA damage response pathways, which is also in agreement with *OPTN* being a highly connected gene to the NetWAS 2.0 module associated with these processes. The other two modules included cytokine secretion, supporting the presence of neuroinflammation, and K48-linked protein ubiquitination. Interestingly, K48 is one of the most common forms of tau ubiquitination for its degradation by the proteasome^18^.

**Figure 1.**
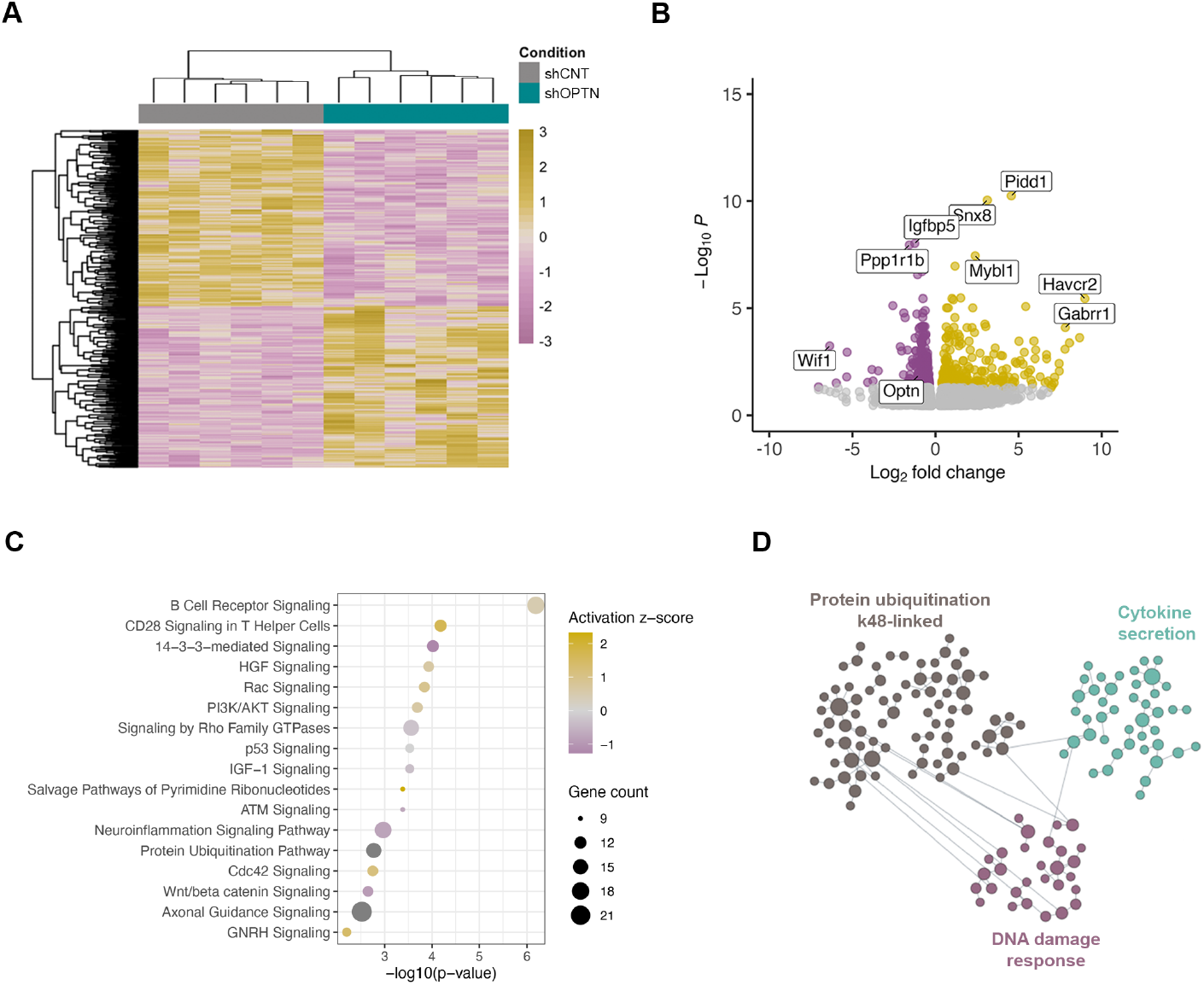
Gene expression changes in ECII neurons triggered by Optn silencing in ECII-neuron bacTRAP mice in vivo. **A)** Heatmap showing the separation between control and Optn-silenced ECII neurons. **B)** Volcano plot showing DEGs between control and Optn-silenced ECII neurons. **C)** Dot-plot showing the most significantly altered pathways in Optn-silenced ECII neurons. **D)** Functional modules identified to be altered in Optn-silenced ECII-neurons 3 week after transduction by HumanBase.

### Proteomics analysis in *Optn*-silenced primary neurons shows early metabolic alterations followed by neuroinflammation and impaired proteostasis

Our data point to a role of optineurin in axon-related processes. Indeed, Optineurin has recently been shown to interact with microtubules and facilitate the transport of mitochondria to axonal terminals, which suggests that its dysfunction may be an essential contributor to axonal degeneration^19^. We thus set out to directly examine alterations in the cytoplasmic and synaptic proteome of *Optn*-silenced neurons *in vitro*. To gain insight into the progression of those changes with time, we analyzed each compartment at 4 and 5 days post transduction (**Figure S2A**). We first verified the downregulation of Optineurin at each time point in whole cell homogenates. Our results showed a significant downregulation of Optineurin only at 5 days post-transduction (**Figure S2B**). We determined the efficient isolation of each compartment by western blot against the nuclear marker Lamin and the pre-synaptic marker SNAP25 (**Figure S2C**). Principal component analysis (PCA) showed a clear separation between the proteomes of each compartment at each time point (**Figure S2D**), which most likely represent changes driven by the ongoing neuronal differentiation at those days in culture. PCA also showed a clear separation between the proteomes of *Optn*-silenced vs. Control neurons, that, in agreement with our western blot analysis of Optineurin levels, were especially apparent at 5 days post-transduction (**Figure S2E**).

Differential protein abundance analysis revealed major alterations in the proteome of both the cytoplasm and synapse at each time point, with increasing changes in the proteome and their significance with time (**Figure 2**). MTHFD2 (methylenetetrahydrofolate dehydrogenase 2) was consistently among the top downregulated proteins at both time points and compartments. MTHFD2 is a mitochondrial enzyme involved in one carbon metabolism from folate, which contributes to mitochondrial complex I function and has been shown to be neuroprotective in the context of amyloid pathology^20^. At 5 days post-transduction we observed a nominally significant decrease in APP in the neuronal cytoplasm (log2FC = -0.2, pval = 0.016), and a nominally significant increase in Tau in the synapse (log2FC = 0.25, pval = 0.024) (**Figure 2F and 2I**). At 4 days post transduction, Ingenuity Pathway Analysis and functional module grouping by HumanBase showed early alterations in mitochondrial function and metabolism in the neuronal cytoplasm. Indeed, in agreement with the observed decrease in MTHFD2, HumanBase identified a module associated with alterations in mitochondrial complex I. Changes in the synapse were less apparent at this time-point, and HumanBase did not find any significant altered functional modules (**Figure 2 A-E**). At 5 days post-transduction, we observed extensive alterations both in the cytoplasm and synaptic compartment with pathways related with proteostasis (including Coordinated Lysosomal Expression and Regulation, CLEAR, in the synapse), endoplasmic reticulum to golgi organization, protein and vesicle transport, synaptic function (including neurexins and neuroligins, especially in the synapse) and neuroinflammation becoming more salient.

**Figure 2.**
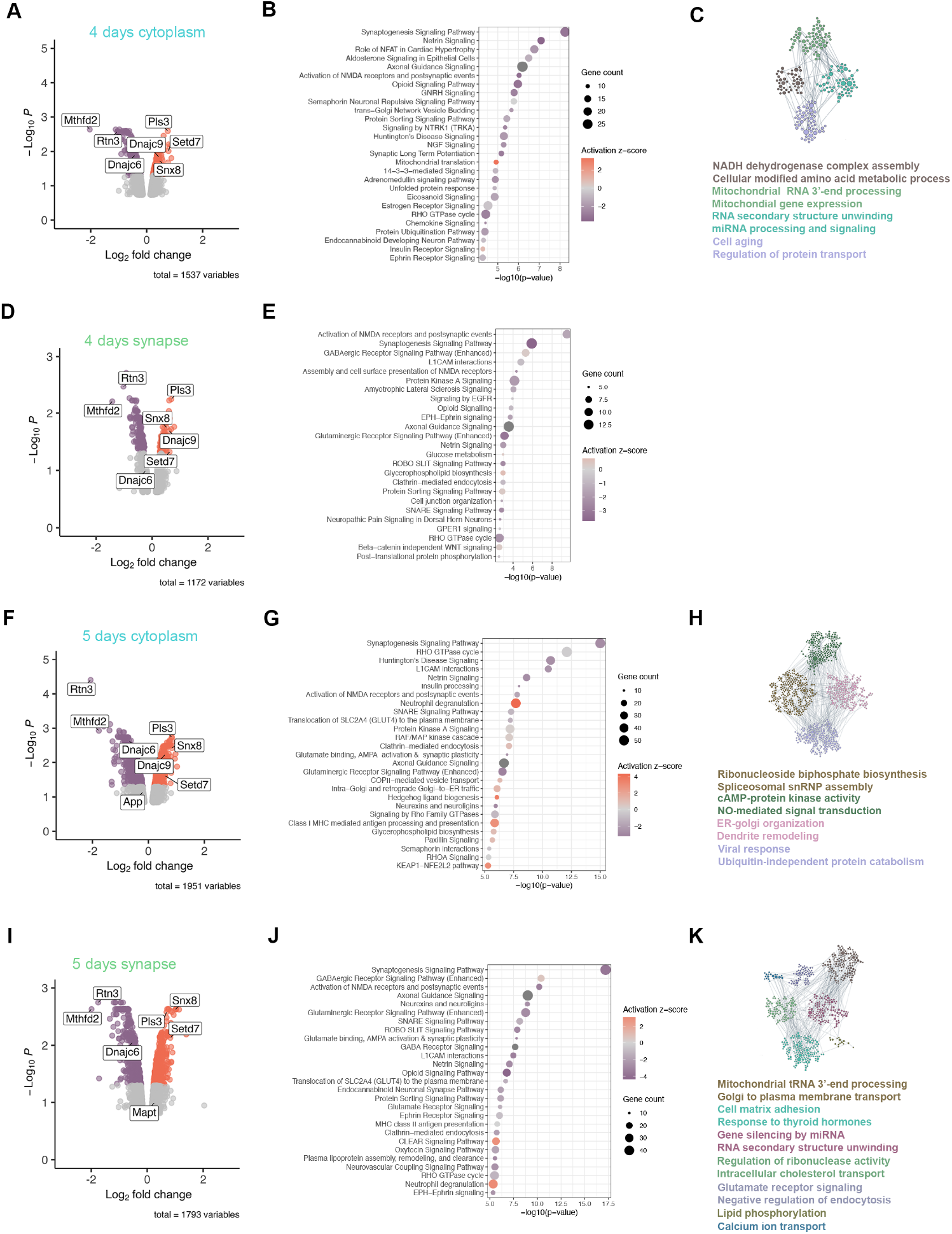
Changes in the subcellular proteome of Optn-silenced neurons in vitro. **A)** Volcano plot of the differentially enriched proteins between control and Optn-silenced cytoplasmic fractions 4 days after transduction. **B)** Dot-plot showing the most significantly altered pathways in the cytoplasm of Optn-silenced neurons 4 days after transduction. **C)** Functional modules identified to be altered in the cytoplasm of Optn-silenced neurons 4 days after transduction by HumanBase. **D)** Volcano plot of the differentially enriched proteins between control and Optn-silenced synaptic fractions 4 days after transduction. **E)** Dot-plot showing the most significantly altered pathways in the synaptic fraction of Optn-silenced neurons 4 days after transduction. **F)** Volcano plot of the differentially enriched proteins between control and Optn-silenced cytoplasmic fractions 5 days after transduction. **G)** Dot-plot showing the most significantly altered pathways in the cytoplasm of Optn-silenced neurons 5 days after transduction. **H)** Functional modules identified to be altered in the cytoplasm of Optn-silenced neurons 5 days after transduction by HumanBase. **I)** Volcano plot of the differentially enriched proteins between control and Optn-silenced synaptic fractions 5 days after transduction. **J)** Dot-plot showing the most significantly altered pathways in the synaptic fraction of Optn-silenced neurons 5 days after transduction. **K)** Functional modules identified to be altered in the synapse of Optn-silenced neurons 5 days after transduction by HumanBase.

We next compared our *in vivo* ECII-neuron bacTRAP data with *in vitro* proteomics to determine genes and proteins that are commonly up or downregulated in *Optn*-silenced neurons (**Figure 3A, 3C**). Pathway analysis of the commonly altered genes and proteins showed alterations in several pathways involved in proteostasis, including protein ubiquitination, cellular response to heat stress or CLEAR signaling pathways. Reelin signaling was also among the top dysregulated pathways (**Figure 3B, 3D**).

**Figure 3.**
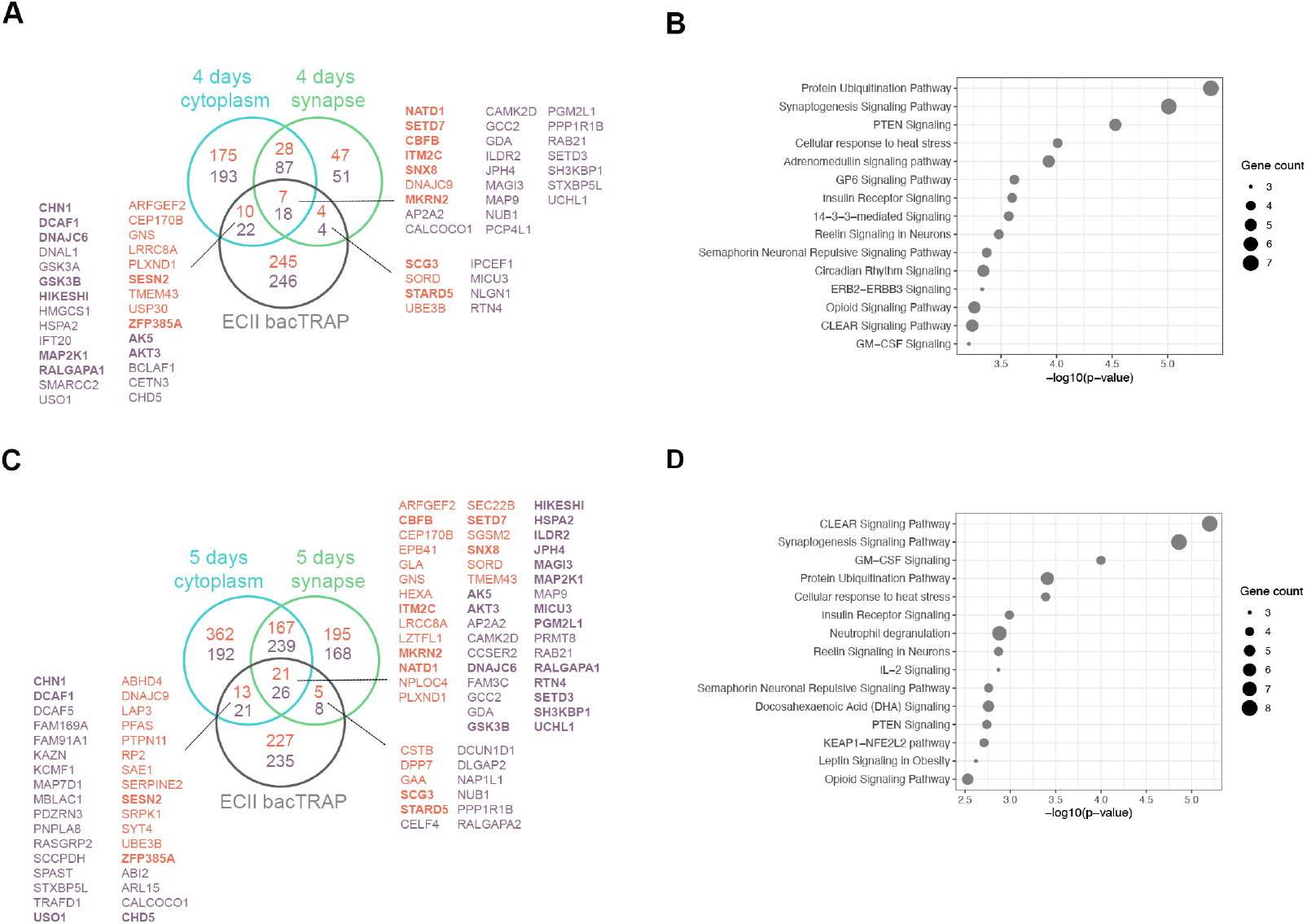
Overlap between DEGs upon Optn silencing in neurons in vivo and differentially enriched proteins in vitro. **A)** Venn diagram showing commonly up (orange) and downregulated (purple) genes and proteins in Optn-silenced neurons from the in vivo bacTRAP data and 4 days post-transduction subcellular in vitro proteomics. **B)** Dot-plot showing the 15 top significant commonly altered pathways in in vivo bacTRAP data and 4 days post-transduction subcellular in vitro proteomics. **C)** Venn diagram showing commonly up (orange) and downregulated (purple) genes and proteins in Optn-silenced neurons from the in vivo bacTRAP data and 5 days post-transduction subcellular in vitro proteomics. Proteins highlighted in bold indicate that they are commonly changed at 4 and 5 days post-transduction. **D)** Dot-plot showing the 15 top significant commonly altered pathways in in vivo bacTRAP data and 5 days post-transduction subcellular in vitro proteomics

### *Optn* downregulation triggers ECII neuron loss and neuroinflammation

To further determine the relevance of Optineurin function for ECII neurons *in vivo*, we performed stereotaxic injections of control or *Optn*-silencing AAVs in the entorhinal cortices of opposite hemispheres of human-tau knock-in mice (hMAPT knock-in), where the entire murine *Mapt* gene is replaced with human *MAPT*. Immunofluorescence analysis at 4 weeks after injection showed high mCherry levels (used as a reporter for viral transduction efficiency), in the EC and in the projecting areas of those neurons in the molecular layer of the dentate gyrus in the control hemisphere, while mCherry signal was absent from EC neuron somas or their projections in the *Optn*-silenced hemisphere (**Figure 4A**). To determine whether this difference was due to ECII neuron loss, we performed immunofluorescence staining for Reelin (used as a marker for ECII neurons) and the neuronal marker NeuN (**Figure 4B**). Our results showed a significant decrease in the percentage of Reelin positive neurons in the *Optn*-silenced hemisphere (**Figure 4C**).

**Figure 4.**
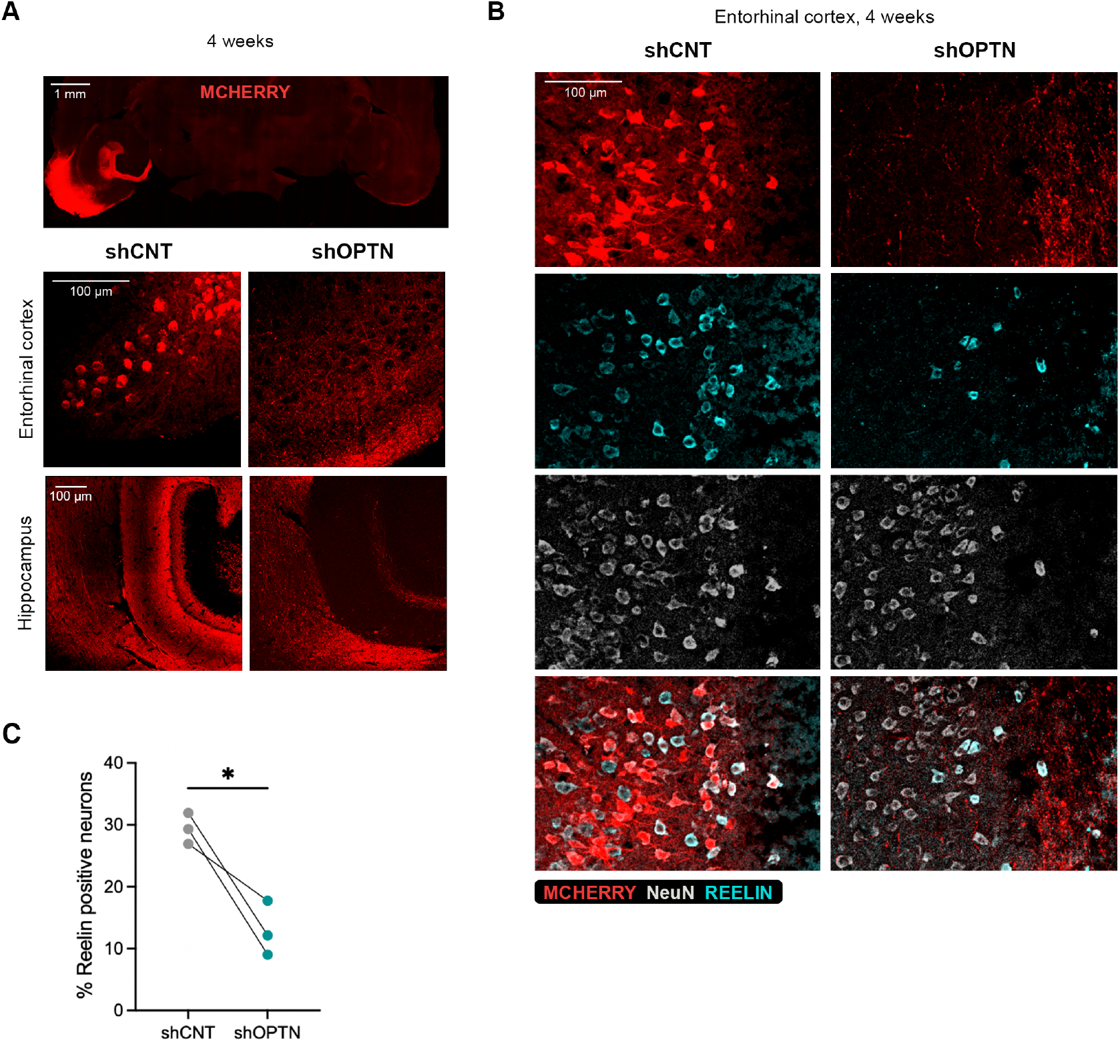
Effect of Optn silencing in ECII neurons in vivo 4 weeks after injections. **A)** Immunofluorescence images of mCherry (red, viral transduction reporter) in the EC and projecting areas of EC neurons in the hippocampus. **B)** Immunofluorescence images of mCherry (red), Reelin (cyan) and NeuN (gray) in the EC. **C)** Graph showing the percentage of Reelin positive neurons relative to the total neuron population (NeuN positive cells) in control or Optn-silenced ECII hemisphere. * pval < 0.05, Paired two tailed t-test.

Considering that pathway analysis showed highly significant alterations in pathways related with neuroinflammation, we next sought to determine whether *Optn*-silencing in ECII neurons leads to glia reactivity. To do so we performed immunofluorescence staining for the microglia marker Iba1 and the astrocytic marker GFAP (glial fibrillary acidic protein). Interestingly, while we did not observe differences in the morphology or abundance of microglia or astrocytes in the EC of the *Optn*-silenced hemisphere at 4 weeks post-injection (**Figure 5A-B**), we did observe a significant increase in GFAP positive signal in projecting areas of EC neurons in the hippocampus (**Figure 5A-B**).

**Figure 5.**
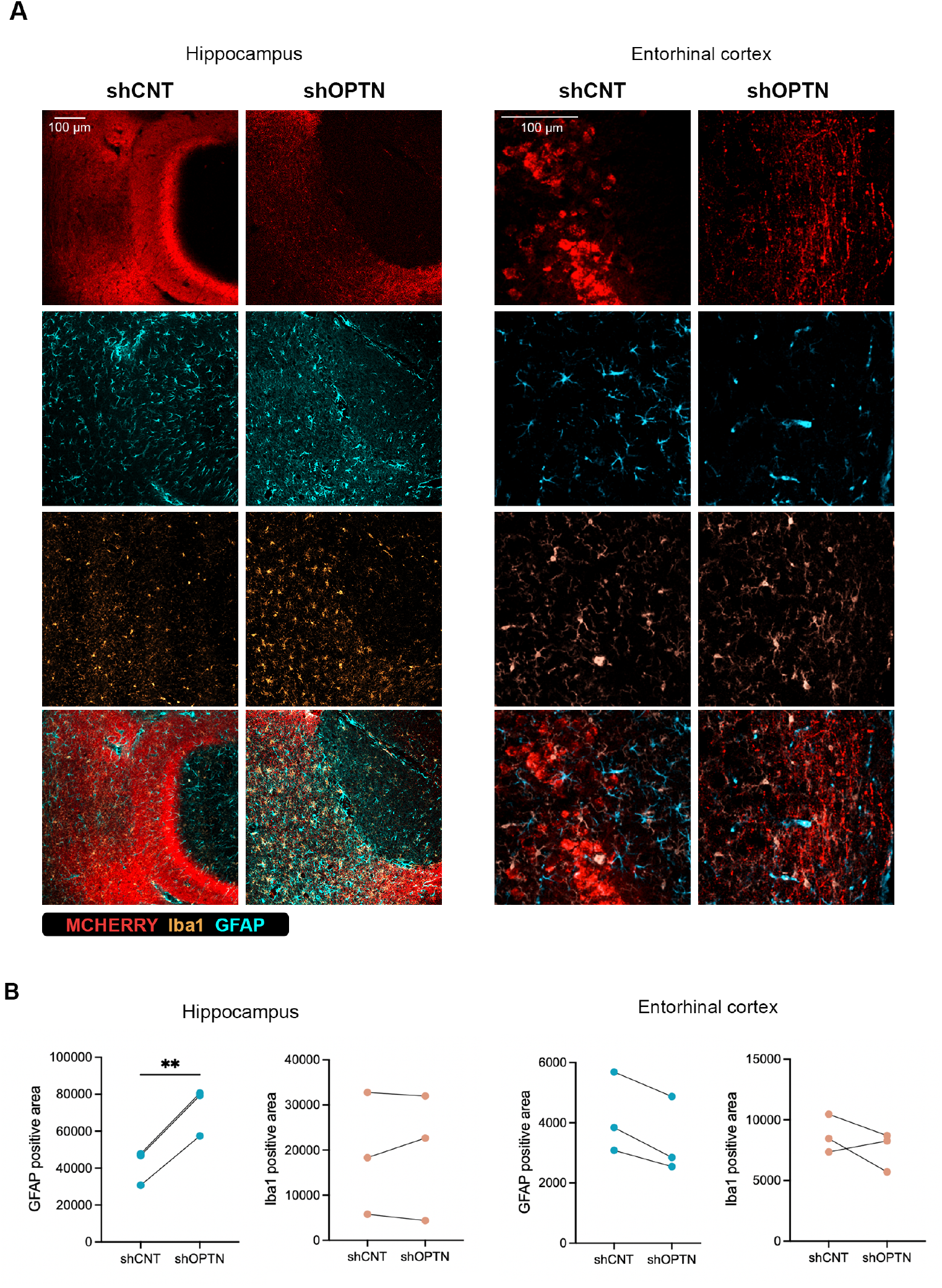
Effect of Optn silencing on ECII neurons in vivo on glial cells 4 weeks after injections. **A)** Immunofluorescence **B)** Graphs showing the abundance of astrocytes (GFAP) or microglia (Iba1) in the hippocampus and EC of control or Optn-silenced hemispheres. ** pval < 0.005, Paired two tailed t-test.

## Discussion

The early degeneration of specific subpopulations of neurons, known as selective neuronal vulnerability, is a common hallmark of neurodegenerative disorders, including AD. These disorders are also commonly characterized by disrupted protein homeostasis that results in the presence of intraneuronal aggregates of certain proteins, such as TDP-43, alpha-synuclein and tau. Despite substantial advances in our understanding of the molecular events that underlie these diseases, the mechanisms that lead to the early degeneration of vulnerable neuron subpopulations remain largely unknown.

To unravel the molecular processes that lead to the early degeneration of ECII neurons in AD, we generated high quality molecular signatures of ECII neurons and other neurons that are resistant to AD throughout the mouse lifespan. We used this information to build ECII neuron-specific functional networks and combined them with AD genetics data to predict potential drivers of tau pathology in AD vulnerable neurons^10^. We previously validated this approach by showing that downregulation of *Dek*, a core regulator of a functional module predicted to drive tau pathology in ECII neurons indeed led to several early AD-associated alterations and tau buildup in vulnerable neurons^21^. *DEK* is, however, not enriched in ECII neurons and therefore whether its effect on Tau is ECII neuron-specific needs to be explored in more detail. In the present work we therefore focus on ECII neuron-enriched genes and identify *Optn*, not only to be highly enriched in these neurons compared to neurons resistant to AD, but to also be a central regulator of a module predicted to drive pathology in AD neurons and that is altered in the presence of amyloid pathology.

Optineurin is known to function as an autophagy adaptor for selective autophagy processes, including mitophagy and aggrephagy, by connecting ubiquitinated cargo to the autophagy machinery through its interaction with LC3^22^. It has also been recently shown to play a central role in the degradation of ubiquitinated substrates through Golgi membrane-associated degradation (GOMED), also known as alternative autophagy^23^. Several mutations in *OPTN* have been associated with ALS and glaucoma and, while its role has been extensively studied in other pathological contexts, its role in AD and in the context of selective neuronal vulnerability has not been explored in detail.

We show that Optineurin downregulation in a non-pathological context (this is, without the presence of amyloid or tau pathology) leads to broad alterations in gene expression and in the subcellular proteome of those neurons. *In vitro* proteomics analysis revealed early alterations in mitochondrial function, especially affecting respiratory complex I. This is followed by progressive impairment in pathways related to proteostasis, synaptic function, and neuroinflammatory signaling. Alterations in mitochondrial function and bioenergetics have long been identified as early events in the pathogenesis of AD^24-26^. However, many of these studies have been performed on pathological contexts where it is the presence of either amyloid or tau pathology that initiates alterations in mitochondrial function, triggering a positive feedback loop that further accelerates oxidative stress, impaired proteostasis, disrupted ATP production, synaptic dysfunction and neurodegeneration^27^. Our results suggest that Optineurin disfunction may act upstream of these events by impairing mitochondrial quality control mechanisms. Consistent with this notion, we observe that selective downregulation of Optineurin in ECII neurons in vivo is sufficient to trigger neuronal loss and astrocyte reactivity in hippocampal projection areas of these neurons, even in the absence of amyloid or tau pathology. These findings suggest that Optineurin dysfunction alone can initiate a cascade of events that are consistent with early stages of AD, including neuronal network disruption and neurodegeneration.

Taken together, our findings position Optineurin as a critical regulator of neuronal homeostasis in AD vulnerable neurons that links impaired mitochondrial quality control with downstream AD-associated alterations. While further work will be needed to investigate the effect of Optineurin downregulation in Tau homeostasis and its potential role in ECII neuron degeneration in our model, it is noteworthy that in a recent high-throughput screen for genes regulating tau oligomer levels in iPSC derived neurons, mitochondrial genes represented the most significant class of gene hits ^28^. *Optn* silencing in ECII neurons might thus lead to some tau proteoform that contributes to ECII neurodegeneration.

Our results also suggest that ECII neurons may be particularly vulnerable to perturbations in mitochondrial homeostasis. The high expression of *Optn* in these cells may reflect an intrinsic neuroprotective mechanism to meet elevated mitochondrial quality control demands that becomes progressively imbalanced by AD-associated stressors (such as amyloid pathology or ageing), rendering these neurons the first to degenerate in this context. Future work will be needed to determine whether restoring either Optineurin function or its downstream targets can protect neurons from degeneration and its translational potential in an AD context.

## Methods

### Mouse models

All experiments involving mouse models were approved by Karolinska Institute (21623-2023) and the Rockefeller University Institutional Animal care and Use Committee (IACUC protocols #16902 and 19067-H) ethical committees.

ECII-bacTRAP mouse are transgenic for the BAC #RP23-307B16 where a cDNA encoding eGFP-L10a (enhanced green fluorescent protein fused to the L10a subunit of the ribosome) was integrated before the start codon of Sh3bgrl2. As a result, they express eGFP-L10a under the control of the regulatory regions of the ECII-enriched Sh3bgrl2 gene, which leads to specific expression of eGFP-L10a expression in ECII neurons. Human-tau knock-in mouse^*29*^ (h*MAPT* knock-in), where the entire murine *Mapt* gene is replaced with human *MAPT* was obtained from the laboratory of Prof. Takomi Saido at Riken. Mice were maintained on a 12h dark/light cycle and provided with food and water *ad libitum*. Experiments were performed in 6 months-old h*MAPT* knock-in mice and 6-8 months-old ECII-bacTRAP mice.

### Mouse primary neuron cultures

Entorhinal or cortico-hippocampal tissue from E17 C57Bl/6J mouse embryos (Jackson, stock# 000664) were incubated at 37°C in 0.05% trypsin/EDTA (#T11493, ThermoFisher) for 10 min. After centrifugation the tissue pellet was dissociated in EBSS containing 0.5 mg/ml DNAse I (#10104159001, Merck) with a glass Pasteur pipette. Cells were seeded at 50,000 cells/cm^2^ in Neurobasal medium (#21103049, ThermoFisher), supplemented with 2% B-27 (#17504044, ThermoFisher) and 2 mM GlutaMAX (#35050061, ThermoFisher), and grown at 37 °C in a humidified 5% CO_2_-containing atmosphere. All experiments were performed after a minimum of 10 days *in vitro* (DIV).

### AAVs

Purified adeno-associated virus (AAVs) stocks were produced by Vector Biolabs. The viruses used for *Optn* silencing were: AAV1-mCherry-U6-mOPTN-shRNA (shRNA sequence: 5’-CCGG-AGCGAGCTGTTCGTGTTGAATCTCGAGATTCAACACGAACAGCTCGCT-TTTTT -3’) and AAV1-mCherry-U6-scrmb-shRNA (control shRNA sequence: 5’-CCGG-CAACAAGATGAAGAGCACCAACTCGAGTTGGTGCTCTTCATCTTGTTG-TTTTT-3’). The viruses used for *Optn* overexpression were: AAV1-hSyn1-mOPTN-IRES-mCherry (expressing the cDNA encoding for mouse OPTN – NM_001356487) and AAV1-hSyn1-mCherry-WPRE (empty control).

### AAVs transduction of mouse primary neurons

Neurons in primary culture at 7 <u>d</u>ays *in <u>v</u>itro* (DIV) were transduced by a full media change to media containing either control, or *Optn*-shRNA AAVs at a concentration of 1 x 10^10^ <u>G</u>enome <u>C</u>opies per 200,000 cells (5 x 10^5^ GC/cell). Neurons were maintained in virus-containing media until the day of the experiment.

### Stereotaxic injections

AAVs were injected in the entorhinal cortex of adult mice using an Angle Two mouse stereotaxic frame with a motorized nanoinjector (Leica) for the experiments performed at The Rockefeller University in ECII-bacTRAP mouse or a 900-model small animal stereotaxic instrument (Kopf) for the experiments performed at Karolinska Institute in h*MAPT* knock-in mice. Animals injected at the Rockefeller University were anesthetized with xylazine (4.5 mg/kg body weight) and ketamine (90 mg/kg body weight) injected intraperitoneally. For the injections performed at the Karolinska Institute, animals were maintained under isoflurane anesthesia (3.5% with 20% oxygen) and supplemented with meloxicam (5 mg/kg) and buprenorphine (0.05 mg/kg) 20 min prior to the surgeries. An ophthalmic ointment was applied to prevent corneal drying during the procedure. AAVs were loaded in a 10 μl syringe (#7653-01, Hamilton) with a 33-gauge needle (#7803-05, Hamilton). 2 μl of AAVs (1 x 10^13^ GC/ml) were injected into the mouse EC (AP: -3.70; ML: -4.65; DV: -4.50) with a nanoinjector tilt of -4°. The same volume and concentration of control AAVs were injected in the contralateral EC (AP: -3.70; ML: +4.65; DV: -4.50) with a nanoinjector tilt of +4°. Injection rate was 0.4 μl/min. Bacitracin antibiotic gel was applied to the surgery wound, that was sutured with a non-absorbable monofilament. To compensate for fluid loss, warm sterile saline solution was injected intraperitoneally (3% of the body weight), and animals were kept on a heated pad and monitored until complete recovery from anesthesia. In cohorts where *Optn* or control AAVs were injected bilaterally in different mice, mice receiving *Optn* or control AAVs were alternated. The experiment unit for the bacTRAP mouse experiments was one mouse (two pooled EC hemispheres). No mouse was excluded from the study after injection.

### bacTRAP-RNAseq

Entorhinal cortices from both hemispheres were isolated from individual mice and processed for bacTRAP as previously described^30^. Tissue was homogenized at 4°C in a glass Teflon homogenizer, in lysis buffer (20 mM Hepes KOH, 10 mM MgCl_2_, 150 mM KCl, 0.5 mM DTT, 100 µg/ml cycloheximide) supplemented with protease inhibitors (#A32965, ThermoFisher) and RNAse inhibitors (40U/ml RNAsin, #N2515, Promega and 20U/ml Superasin, #AM2696, ThermoFisher). After centrifugation, the supernatant was incubated with 1% NP-40 and 30 mM DHPC (#850306P, Avanti) on ice for 5 min. The supernatant, containing ribosome bound RNAs, was then incubated overnight with magnetic beads (Streptavidin MyOne T1 Dynabeads, #65602, ThermoFisher) previously coated with anti EGFP antibodies for immunoprecipitation (HtzGFP-19C8 and HtzGFP-19F7, from MSKCC monoclonal antibody facility ^17^). Immunoprecipitated RNAs were then purified using the RNeasy Plus Micro Kit (#74034, Qiagen). RNA integrity was determined with a Bioanalyzer 2100 (Agilent) using an RNA 6000 pico chip (#5067-1513, Agilent). RNA was quantified with Quant-it Ribogreen RNA reagent (#R11490, ThermoFisher). Reverse transcription was performed with Ovation RNAseq v2 kit (#7102, NuGEN) from 5 ng of RNA following the manufacturer’s instructions. cDNAs were purified using the QIAquick PCR purification kit (#28104, Qiagen). cDNA yield was measured with Quant-IT Picogreen dsDNA kit (#P7581, ThermoFisher). 200 ng of cDNA were used for fragmentation prior to cDNA library preparation. cDNA was sonicated into 200 bp fragments using a Covaris S2 ultrasonicator instrument (10% duty cycle, intensity 5, 200 cycles/burst per second for 2 min at 5.5°C to 6°C). Library preparation was performed with the TruSeq RNA sample preparation kit v2 (#RS-122-2001, Illumina) and were sequenced at the Rockefeller University genomics resource center on a NextSeq 500 sequencer (Illumina).

### Differential gene expression analysis

Following sequencing, adapter and low-quality bases were trimmed by fastp^31^ from the raw sequencing files in FASTQ format. Cleaned reads were aligned to the mm10 reference genome using STAR version 2.7.1a^32^. After alignment, the Reads Per Kilobase of transcript per Million mapped reads (RPKM) for all genes in each sample were calculated with R package edgeR^33^. To analyze differential gene expression between samples, DESeq2^34^ was used, applying the standard comparison mode between two experimental groups.

### Mouse brain cryosections

Mice were anesthetized with isoflurane and perfused with PBS followed by 4% paraformaldehyde (PFA). Brains were then dissected and post-fixed in 4% PFA for 1 h. Afterwards they were washed with PBS and incubated in increasing sucrose concentrations (5%, 15% and 30%) for cryopreservation. They were then embedded in OCT compound (TissueTek), cut in 20 μm-thick free-floating horizontal sections with a CM1860 Cryostat (Leica) and preserved in cryoprotectant solution (30% ethylene glycol, 40% glycerol in PBS) at -20°C until further analysis.

### Subcellular fractionation

Synaptosome and cytosolic fractions from neurons in primary culture were generated with the Syn-PER^TM^ Synaptic Protein Extraction reagent kit (#87793, ThermoFisher), following the manufacturer’s instructions. Briefly, primary neurons were washed in PBS and lysed in Syn-PER^TM^ for ten minutes on ice. Lysates were then centrifuged at 1200 xg for 10 min at 4°C. The pellet (P1) was resuspended in 70 µl RIPA buffer and centrifuged at 10000 xg for 10 min at 4°C, while the supernatant (S1) was centrifuged at 15000 xg for 20 min at 4°C. The resulting supernatants (S2) were combined and constitute the cytoplasmic fraction, while the resulting pellet from S1 (P2) was resuspended in 60 µl Syn-PER^TM^ and constitutes the synaptosomal fraction. All buffers were supplemented with 10% protease (#A32965, ThermoFisher) and phosphatase (#4906837001, PhoSTOP, Merck) inhibitors.

### Western blot

For whole cell homogenates, cells were homogenized in RIPA buffer (#89901, ThermoFisher), supplemented with 10% protease (#A32965, ThermoFisher), and phosphatase (#4906837001, PhoSTOP, Merck) inhibitors. Whole cell extracts or subcellular fractions (cytoplasm and synaptosomes) were then subjected to sodium dodecyl sulfate (SDS) polyacrylamide gel electrophoresis and transferred to a nitrocellulose membrane. Membranes were blocked in Intercept (TBS) blocking buffer (#927-60001, Li-Cor) 1h at room temperature and blotted with the primary antibodies overnight at 4 °C. Antibodies used were Optineurin (1:500, #ab213556,Abcam), SNAP25 (1:500, #ab108990, Abcam), Lamin (1:5000, #SAB4200236, Sigma), Tau (1:1000, #A0024, Dako) and phospho-tau Ser202, Thr205 (AT8, 1:2000, #MN1020, Invitrogen). Actin (1:1000, #A2066, Sigma) was used as loading control. The day after membranes were incubated with IRDye 800CW and 680RD mouse and rabbit secondary antibodies (Li-cor) for 1h at room temperature. Signal detection was performed with an Odyssey scanning system (Li-Cor). Band intensity quantification was performed with the Image Studio Lite software.

### Immunofluorescence

20 μm-thick free-floating horizontal sections were permeabilized in 2% normal donkey serum and 0.1% Triton X-100 in PBS for 30 min at room temperature. They were then stained overnight at 4°C with primary antibodies diluted in the same permeabilization buffer. Primary antibodies used were mCherry (1:1000, #ab205402, Abcam), Reelin (1:50, #PA5-47537, Invitrogen), NeuN (1:500, #ABN78, Merck), Iba1 (1:500, #019-19741, Wako), GFAP (1:500, #556330, BD) and phospho-tau Thr231 (1:500, #NB100-82249, Novus Biologicals). The day after samples were washed in PBS and incubated with secondary antibodies for 1h at room temperature. The secondary antibodies used were Donkey anti-goat IgG (H+L) Cross-adsorbed Alexa Fluor Plus 405 (#A48259, ThermoFisher), Donkey anti-chicken IgY (H+L) Cross-adsorbed Alexa Fluor 594 (#A78951, ThermoFisher) and Donkey anti-mouse IgG (H+L) Cross-adsorbed Alexa Fluor 647 (#A31571, ThermoFisher), Donkey anti-mouse IgG (H+L) Cross-adsorbed Alexa Fluor 488 (#A21202, ThermoFisher) and Donkey anti-rabbit IgG (H+L) Cross-adsorbed Alexa Fluor Plus 647 (#A32795, ThermoFisher). After washing with PBS autofluorescence was quenched with Eliminator reagent (#2160, Millipore Sigma) following the manufacturer’s instructions and mounted with ProLong Gold Antifade Reagent (#P36980, ThermoFisher).

### Image acquisition

Images were acquired on a Zeiss Axio Scan.Z1 (Carl Zeiss) automated high content imaging system or on Zeiss LSM900-airy or Zeiss LSM800-airy laser scanning confocal microscopes (Carl Zeiss).

### Immunofluorescence images quantifications

Image quantification of Reelin positive neuron abundance was performed using QuPath 0.6. EC layer II was selected as the area of interest for each hemisphere and a minimum of 3 sections per mouse (n = 3 injected mice). Cell segmentation was performed using the NeuN signal to select neurons within that area. Reelin positiveness was selected using supervised thresholding. All the cells that showed a higher mean signal than the threshold were considered positive. This allowed us to generate a classification of Reelin +/-neurons and calculate their percentage over the total neuron population in ECII. The averaged percentage for each mouse was used for statistical analysis.

Image quantification of astrocyte and microglia abundance was performed in QuPath 0.6 with the pixel classification feature and supervised thresholding. The total area within a region of interest that was classified above the threshold was used for statistical comparisons between control and *Optn*-silenced hemispheres.

### Mass spectrometry analysis

Mass spectrometry analysis was performed by the Clinical Proteomics Mass Spectrometry facility, Karolinska Institute/ Karolinska University Hospital/ Science for Life Laboratory. Briefly, synaptosome and cytosolic fractions from *Optn*-silenced or control neurons at 4 and 5 days post-transduction were further lysed in 4 % SDS and digested using a modified version of the SP3 protein clean up and digestion protocol^35^ followed by SP3 peptide clean up procedure. Peptide separation was performed on an Evosep One system (Evosep, Denmark) coupled to a timsTOF HT mass spectrometer (Bruker Daltonics, Germany). Samples were run using the 30SPD (44-minute) gradient in DIA-PASEF acquisition mode. Acquired data was analyzed using Spectronaut in Direct-dia mode, with Mus Musculus (Mouse) Uniprot database for direct database matching.

### Differential protein enrichment analysis

Mass spectrometry data was analyzed using R 4.4.3/Rstudio (2026.01.0). Protein accession numbers were mapped to gene names and functional descriptions using Ensembl BioMart (*Mus Musculus*), retrieving UniProt Swiss-Prot identifiers, gene names, and annotation metadata for downstream analysis. Proteins with fewer than two precursors in more than 25% of the samples were excluded from the dataset. All proteins identified as keratins were excluded from the dataset as potential contaminants. MS1 values were log2-transformed and normalized to the median. Statistics were done using the limma package^36^, and p-values were corrected using the Benjamini-Hochberg method. Only proteins with an adjusted p-value lower than 0.05 were considered differentially enriched.

### Statistical analysis

Statistical analyses were performed with GraphPad Prism 10. Statistical details can be found in the figure legends. On the figures throughout the manuscript, we use the usual p-value convention: 0.01 < p-value ≤ 0.05: *; 0.001 < p-value ≤ 0.01: **; 0.0001 < p-value ≤ 0.001 ***; 0.0001 ≤ p-value: ****

## Abbreviations

AD: Alzheimer’s Disease
AAVs: Adeno-associated virus
bacTRAP: bacterial artificial chromosome – Translating Ribosome Affinity Purification
DEGs: differentially expressed genes
DGE: differential gene expression
EC: entorhinal cortex
DIV: days in vitro
ECII: entorhinal cortex layer II
eGFP: enhanced green fluorescent protein
GC: genome copies
IEGs: Immediate Early Genes
NetWAS: Network-Wide Association Study
NFTs: Neurofibrillary tangles.

## Author contribution

Authors contribution is assigned following CRediT (Contributor Roles Taxonomy) guidelines. Conceptualization and methodology, P.R-R. and J-P.R.; Formal Analysis, W.W., C.T., J.S.W., M.T, J-P.R. and P.R-R.; Investigation, J-P.R., C.T., J. S.W., M.T., C.G-M., Y.T., B.M., M.L-L. and P. R-R.; Resources, P.R-R. and J.P.R.; Writing-Original Draft, P.R-R. and J-P.R.; Writing-Review & Editing, C.T., J.S.W. and M.T.; Visualization, P.R-R., C.T. and J.S.W.; Funding Acquisition, P.R-R and J.P.R.

## Acknowledgements

This work was supported by the European Union’s Horizon 2020 research and innovation program under the Marie Sklodowska-Curie grant agreement No 799638 (to P.R-R.); the Fisher Center for Alzheimer’s Disease Research (to J-P.R. and P.R-R.); Cure Alzheimer’s Fund (to J-P.R.); Alzheimerfonden (to P.R-R.); the Karolinska Institute fund for Doctoral education (KID) (to P.R-R.); The private initiative “Innovative ways to fight Alzheimer’s disease-Leif Lundblad family and others” (to P.R-R.); Petrus and Augusta Hedlunds foundation (to P.R-R.); Demensfonden (to P.R-R.); Konung Gustaf V:s och Drottning Victoria foundation (to P.R-R.); Gamla Tjänarinnor foundation (to P.R-R. and C.T.); Gun and Bertil Stohnes foundation (to P.R-R. and C.T.); the National Institute on aging of the NIH (awards RF1 AG047779 and R21 AG085464) (to J-P.R.); Alzheimer’s Association (AARGD-22-932597) (to J-P.R.) and grants from the National Center for Advancing Translational Sciences (NCATS, National Institutes of Health (NIH) Clinical and Translational Science Award) #UL1TR001866 (CTSA) program and BU-CTSI #1UL1TR001430 (to J-P.R.). Its contents are solely the responsibility of the authors and do not necessarily represent the official views of the NIH.

We thank C. Zhao, C. Lai and the whole Rockefeller University genomics resource center team for all the sequencing. We thank T. Carroll and the Rockefeller University Bioinformatics facility for their help with data analysis. We also thank the Clinical Proteomics Mass Spectrometry Facility at Science for Life Laboratory for the mass spectrometry analysis. We thank Per Nilsson for his work to make the h*MAPT* knock in mouse line available at our department at the Karolinska Institute, and Cecilia Dominguez for coordinating the breeding of the mouse cohorts necessary for this study.

## Data availability

The mass spectrometry proteomics data have been deposited to the ProteomeXchange Consortium^37^ via the PRIDE^38^ partner repository with the dataset identifier PXD075358.

## Extended data figures

**Supplementary Figure 1.**
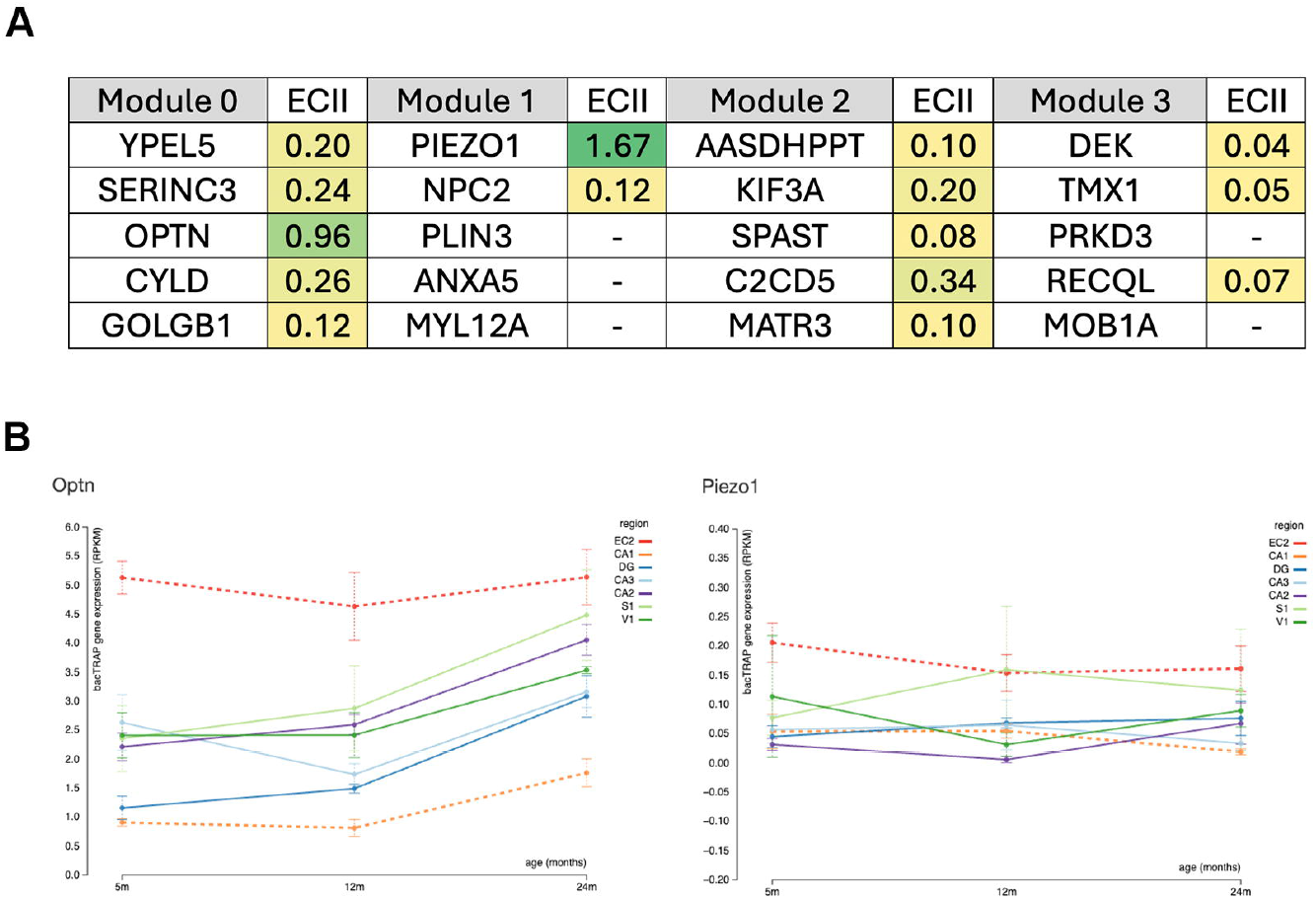
**A)** Top 5 genes connected to each NetWAS 2.0. module and their enrichment score in ECII neurons (ECII) compared to neurons resilient to AD. **B)** Gene expression levels of *Optn* and *Piezo1* throughout the mouse lifespan in different brain neurons from bacTRAP data. Source: alz.princeton.edu.

**Supplementary Figure 2.**
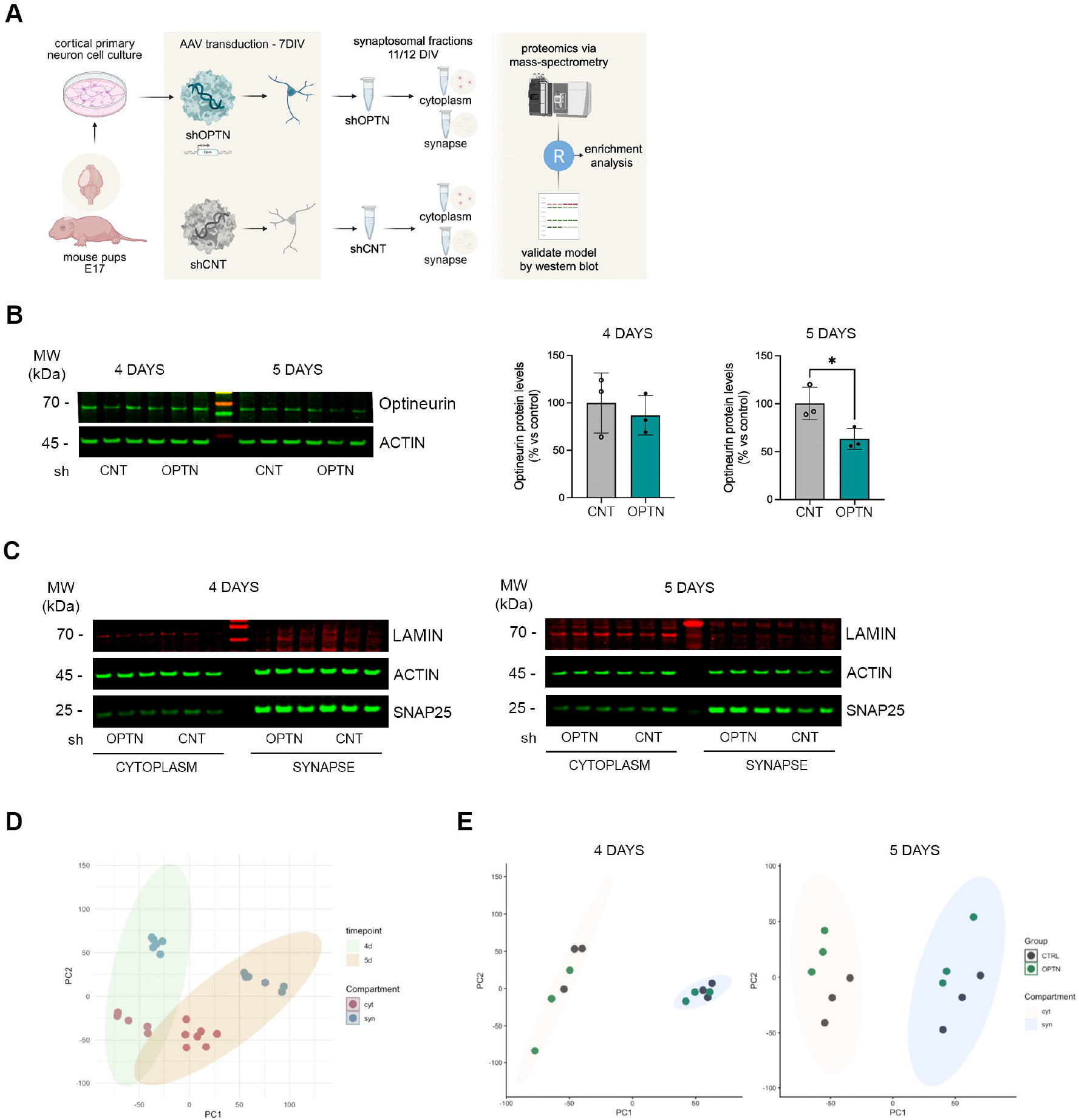
**A)** Schematic representation of the *in vitro* proteomics experiments. **B)** Western blots showing Optineurin levels at 4 and 5 days post-transduction in whole cell homogenates. **C)** Western blots showing efficient subcellular fractionation of cytoplasm and synaptic compartments. **D)** PCA plot showing the separation between subcellular compartments at 4 and 5 days post-transduction. **E)** PCA plots showing the separation between control and *Optn*-silenced conditions for each of the compartments at 4 and 5 days post transduction. * pval < 0.05, Unpaired t-test with Welch’s correction.

**Supplementary Table 1**. Gene expression changes in ECII neurons triggered by up and downregulation of *Optn in vivo*.

**Supplementary Table 2**. Pathways altered in *Optn*-silenced ECII neurons *in vivo* according to ingenuity pathway analysis.

**Supplementary Table 3**. Subcellular protein abundance changes triggered by *Optn* downregulation in neurons in primary culture.

**Supplementary Table 4**. Pathways altered in *Optn*-silenced neuron cytoplasm and synapse *in vitro* according to ingenuity pathway analysis.

